# GFFx: A Rust-based suite of utilities for ultra-fast genomic feature extraction

**DOI:** 10.1101/2025.08.08.669426

**Authors:** Baohua Chen, Dongya Wu, Guojie Zhang

**Affiliations:** School of Basic Medical Sciences, Zhejiang University School of Medicine, Hangzhou 311121, China; Center for Evolutionary & Organismal Biology, Liangzhu Laboratory, Zhejiang University Medical Center, Hangzhou 311121, China

**Keywords:** GFF file, Genome Annotation, Rust Programming, Feature Extraction

## Abstract

Genome annotations are becoming increasingly comprehensive due to the discovery of diverse regulatory elements and transcript variants. However, this improvement in annotation resolution poses major challenges for efficient querying, especially across large genomes and pangenomes. Existing tools often exhibit performance bottlenecks when handling large-scale genome annotation files, particularly for region-based queries and hierarchical model extraction. Here, we present *GFFx*, a Rust-based toolkit for ultra-fast and scalable genome annotation access. *GFFx* introduces a compact, model-aware indexing system inspired by binning strategies and leverages Rust’s strengths in execution speed, memory safety, and multithreading. It supports both feature- and region-based extraction with significant improvements in runtime and scalability over existing tools. Distributed via Cargo, *GFFx* provides a cross-platform command-line interface and a reusable library with a clean API, enabling seamless integration into custom pipelines. Benchmark results demonstrate that *GFFx* offers substantial speedups and makes a practical, extensible solution for genome annotation workflows.

## Introduction

With the growing understanding of functional genome regions beyond conventional protein-coding genes, genome annotations are rapidly increasing in both complexity and volume. Large-scale efforts such as ENCODE [1], FANTOM [2], and Roadmap Epigenomics Program [3] have cataloged diverse noncoding elements—including enhancers, promoters, long non-coding RNAs (lncRNAs), and epigenetic marks—highlighting their roles in gene regulation, chromatin dynamics, and cellular identity. As novel regulatory elements, alternative isoforms, and lineage- or tissue-specific transcripts continue to emerge, annotation datasets are expected to expand further [4]. The accumulation of such multilayered annotations, particularly across large genomes or pangenomes, poses growing challenges for storage, indexing, and efficient querying.

However, existing tools often struggle to process ultra-large annotation files efficiently, particularly for region-based queries, hierarchical model extraction, or parallel execution. A scalable, high-performance solution optimized for such tasks is urgently needed. Rust, a modern systems programming language, offers high execution speed, memory safety, efficient multithreading, and cross-platform portability. These features have led to its increasing adoption in bioinformatics [5], as exemplified by Rust-Bio [6], Bigtools [7], Phylo-rs [8], and fibertools [9].

To address these challenges, we developed *GFFx*, a Rust-based toolkit for fast and scalable access to genome annotation files. *GFFx* fills a gap in current methods by enabling efficient indexing and querying of ultra-large General Feature Format (GFF) datasets. Designed as both a command-line tool and a reusable library, it can be integrated into larger pipelines and software systems. It also demonstrates Rust’s potential in computational biology by providing a robust, extensible foundation for high-performance annotation processing.

## Findings

### Performance benchmark in annotation indexing

*GFFx* achieves high-performance efficiency through a modular indexing system anchored by two core indices, .prt and .gof, which capture feature hierarchical relationship and map annotation blocks to their byte-offsets for direct memory access, respectively. Complementary lightweight indices, including .*fts*, .*a2f*, .*atn*, .*sqs, rit*, and .*rix*, support subcommand-specific operations like feature extraction, attribute-based searches, and region queries with minimal I/O overhead (**Fig. 1**).

**Figure 1.**
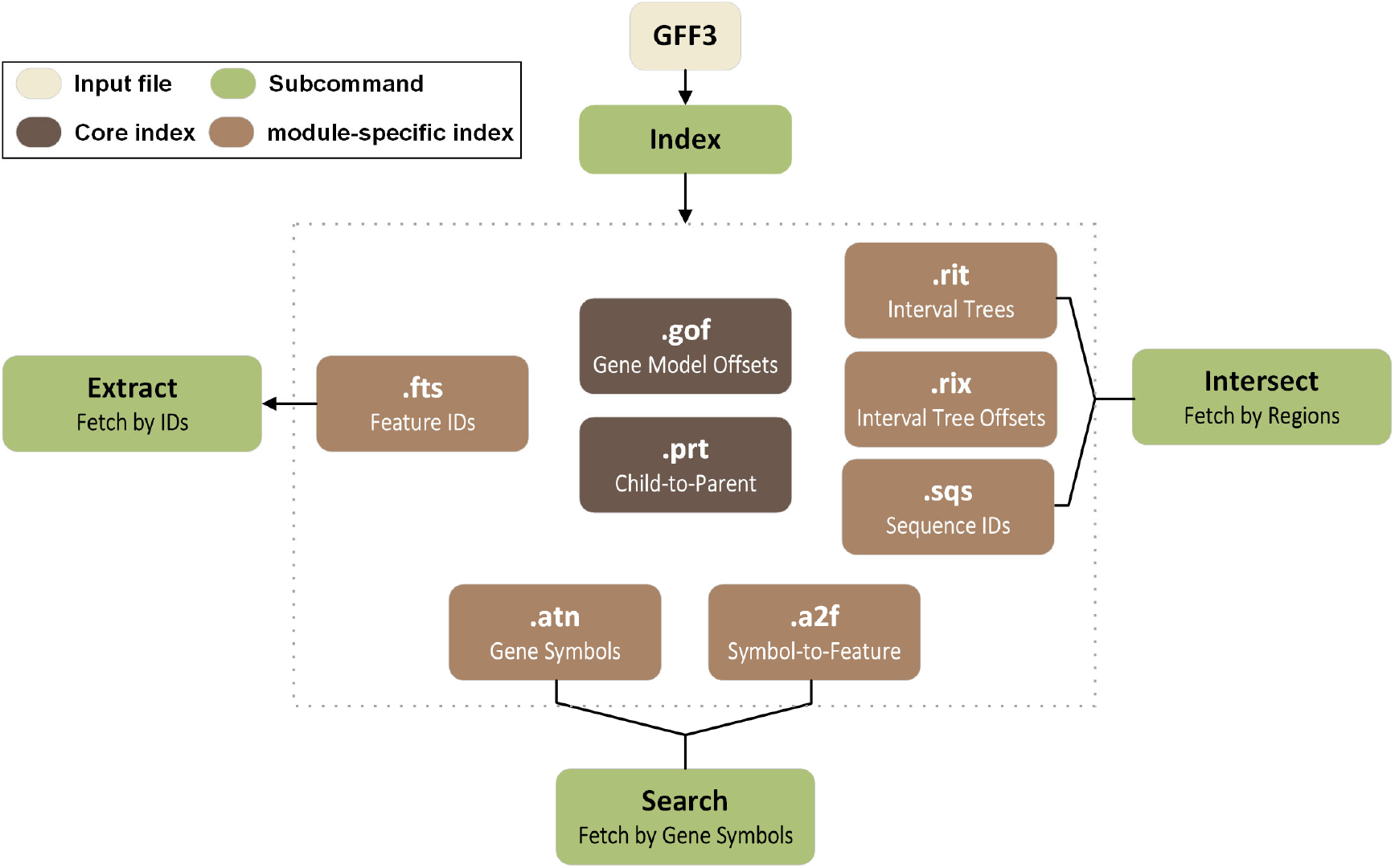
Architecture of the indexing system and subcommand interactions in *GFFx*. All index files are generated in advance from a GFF3 file (cream box) via the index module. While all subcommands (green boxes) have access to the complete set of indices, each subcommand loads only the subset relevant to its specific function. Core indices .gof and .prt (dark brown boxes) are universally required, whereas module-specific indices such as .fts, .a2f, .atn, .rit, rix, and .sqs (light brown boxes) are utilized only by specific subcommands as illustrated.

Among commonly used GFF processing tools, only *gffutils* [10] preprocesses on GFF files by converting it into a SQLite database. We compared the runtime of index-building in *GFFx* and database-creating in *gffutils* (v0.13) using four representative GFF3 annotation datasets: the *Homo sapiens* genome (hg38), the *Pungitius sinensis* genome (ceob_ps_1.0), the *Drosophila melanogaster* genome (dm6), and the *Arabidopsis thaliana* genome (tair10.1), which contain 4 894 761, 652 095, 412 995 and 710 661 feature items, respectively. All benchmarks were performed on a dedicated compute node equipped with 2 × Intel(R) Xeon(R) Gold 6448H CPUs (32 cores/64 threads each), 1 TB DDR4 RAM, and dual Micron 7450 MTFDKCB960TFR NVMe SSDs (total capacity 1.92 TB). The results indicate that, despite its relatively complex indexing architecture, *GFFx* did not exhibit a significant increase in runtime. For GFF files of moderate size, *GFFx* took one to two minutes—approximately 1.44 to 2.17 times longer than *gffutils*. However, for the large hg38 GFF file, *GFFx* outperformed *gffutils* substantially, running 6.72 times faster (**Supplementary Fig. S1**). In addition, the total sizes of the index files generated by *GFFx* were remarkably smaller than those of database files produced by *gffutils* (**Supplementary Table S1**), underscoring another key advantage of *GFFx*.

### Benchmarking identifier-based feature extraction performance

We benchmarked identifier-based feature extraction performance of *GFFx* against four existing tools: *gffread* (v0.12.8) [11], *gffutils* (v0.13) [10], *bcbio-gff* (v0.7.1) [12] and *AGAT* (v1.4.1) [13]. These benchmarks used the same four annotation GFF files as above, with 100 replicates per file. In each replicate, we randomly sampled 100,000 feature identifiers and extracted the corresponding entries. *GFFx* achieved median runtimes of 1.62 s, 0.41 s, 0.37 s and 0.38 s on hg38, ceob_ps_1.0, dm6 and tair10.1, respectively (**Fig. 2a**; **Supplementary Table S2**), corresponding to 12.33-to 80.27-fold speedups over the second fastest tool, *gffread*. Besides, *GFFx* required less memory than other tools except *gffutils* (**Fig. 2b**; **Supplementary Table S2**). Overall, *GFFx* achieves substantial speedups, with the speed increasing proportionally with the size and complexity of the annotation files, without incurring additional memory overhead. As genome assemblies become larger and the annotations grow more detailed, *GFFx* will continue to outpace other tools by an ever-widening margin.

**Figure 2.**
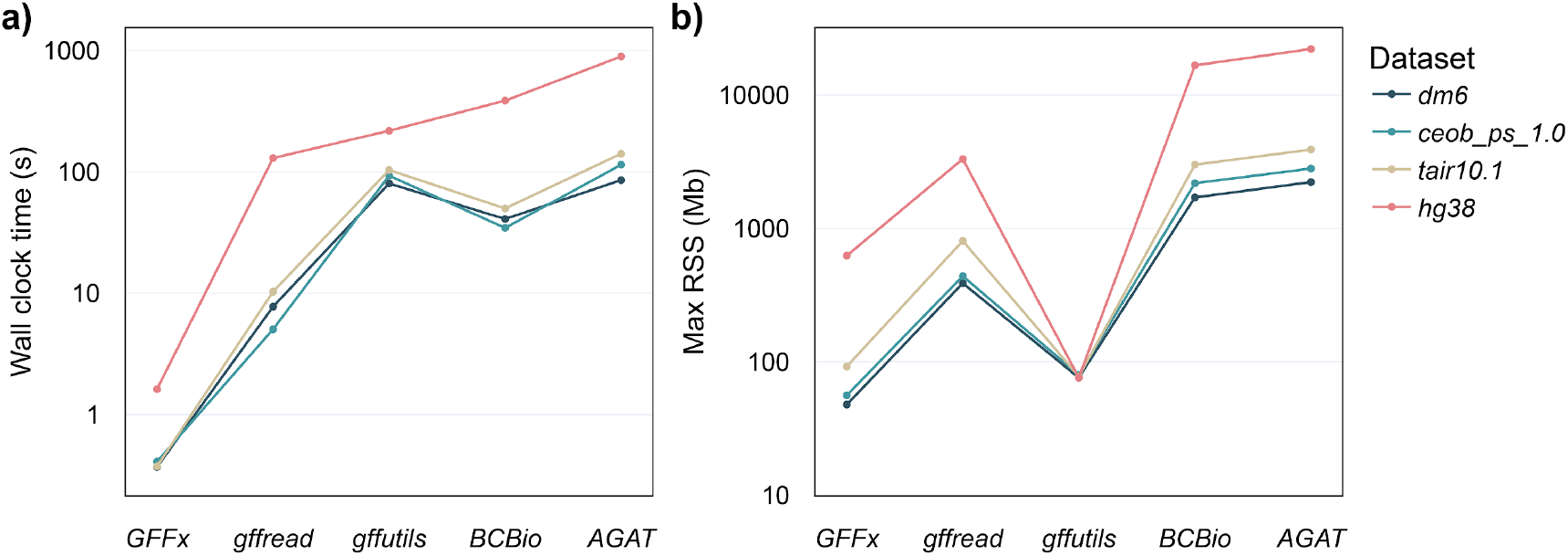
Comparison of identifier-based extraction performance among *GFFx* and other tools. **(a)** Median wall-clock time (log scale) for extracting 100 000 random 20-kbp intervals in dm6 (dark teal), ceob_ps_1.0 (cyan), tair10.1 (tan) and hg38 (pink) using *GFFx, gffread, gffutils, BCBio*, and *AGAT*. **(b)** Maximum resident set size (RSS, log scale), a measure of peak memory consumption, for each tool and dataset. Data represent the median of 100 replicate runs. Tools are ordered left-to-right by increasing median wall-clock time on hg38.

### Benchmarking region-based feature retrieval performance

Subsequently, we benchmarked region-based retrieval performance of *GFFx* against four tools— *gffutils, bcbio-gff, AGAT* and *bedtools* (v2.31.1) [14]—substituting *bedtools* for *gffread* because *gffread* only handles single user-specified regions and does not accept BED files. Using the same four annotation GFF files with 100 replicates each, we generated BED4-format interval files containing 100,000 randomly sampled 20-kbp bins per replicate using the random command from bedtools, which generates random genomic intervals. Among all tools, *GFFx* delivered the fastest region-based retrieval, with median runtimes of 0.46 s, 0.10 s, 0.18 s and 0.17 s on hg38, ce11_ps_1.0, dm6 and tair10.1, respectively (**Fig. 3a**; **Supplementary Table S3**). Excluding *GFFx, bedtools* was the next fastest, requiring 4.89–11.04 s (24.0–61.8-fold slower), while dedicated GFF processors were at least 201-fold slower. This performance gain of *GFFx* derives from its interval-tree algorithm, which reduces time complexity from O(*N*) to O(log *N* + *k*), where *N* represents the total number of intervals in a GFF file and *k* represents the number of overlapped intervals. Although the memory usage of *GFFx* is not always the lowest (**Fig. 3b**; **Supplementary Tables S3**), it remains under 1 GB across all tests, ensuring operability on standard personal computers without sacrificing speed.

**Figure 3.**
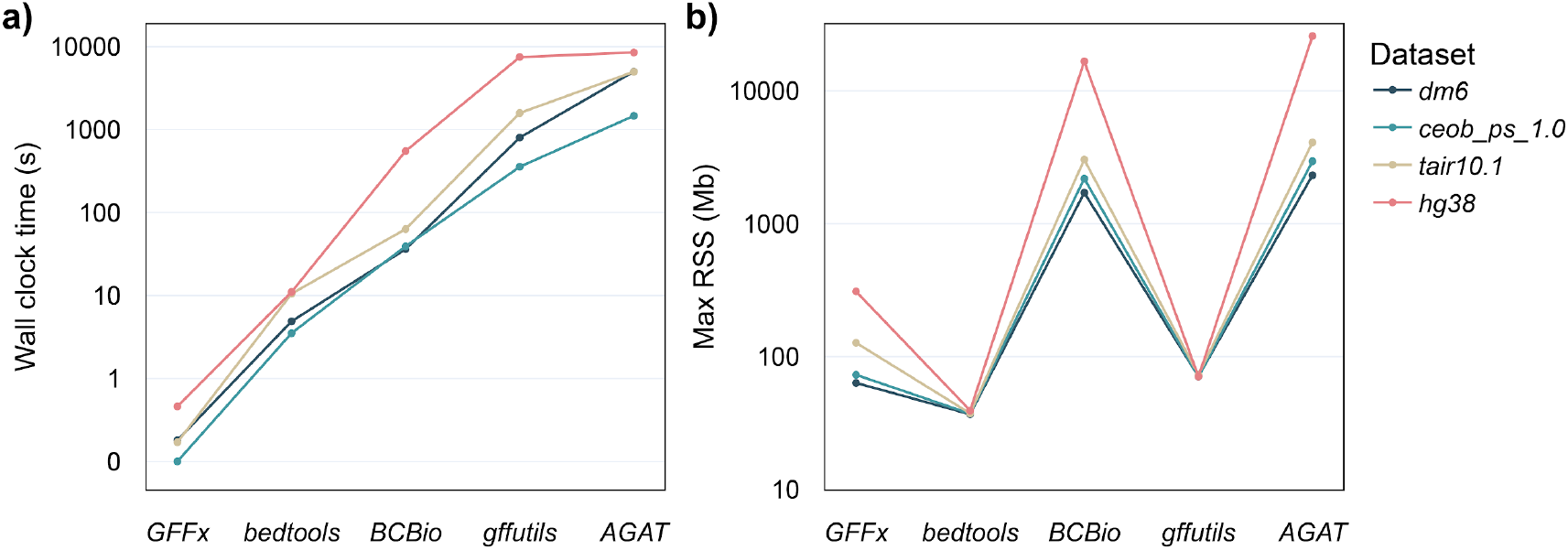
Comparison of region-based feature retrieval performance among *GFFx* and other tools. **(a)** Median wall-clock time (log scale) for extracting 100 000 random 20-kbp intervals in *dm6* (dark teal), *ceob_ps_1*.*0* (cyan), *tair10*.*1* (tan) and *hg38* (pink) using *GFFx, bedtools, BCBio, gffutils* and *AGAT*. **(b)** Maximum resident set size (RSS, log scale), a measure of peak memory consumption, for each tool and dataset. Data represent the median of 100 replicate runs. Tools are ordered left-to-right by increasing median wall-clock time on hg38.

## Discussion

Here, we present *GFFx*, a Rust-based, modular, and high-performance toolkit for efficient processing and querying of ultra-large GFF3 genome annotation files. It addresses key limitations of existing tools through a compact, model-aware indexing system and by leveraging Rust’s strengths in speed, memory safety, and multithreaded execution. Many widely used tools suffer from performance bottlenecks when processing large-scale annotations. For example, *gffutils* depends on relational databases, leading to long indexing times and high disk usage; *AGAT* and *bcbio-gff* offer broad functionality but are not optimized for fast querying; *bedtools* supports region-based queries but lacks model awareness; and *gffread* performs well only on small datasets and lacks parallel support.

Region-based queries in *GFFx* are powered by an in-memory interval-tree index. Interval trees are a well-established data structure for efficiently storing and querying one-dimensional intervals that vary widely in length and often overlap or nest, making them an ideal fit for genome annotation data [15]. In an interval tree, each node represents a feature interval and tracks the maximum endpoint of its subtree. This pruning mechanism skips entire subtrees whose intervals lie outside the query region, avoiding full-file scans and enabling sublinear query times. Once features are identified, *GFFx* uses the .*gof* index, which maps feature IDs to byte offsets in the original GFF file, to retrieve annotation blocks directly, resulting in rapid end-to-end extraction even on large, complex datasets.

Benchmark results show that *GFFx* significantly outperforms existing tools in both feature- and region-based extraction, offering large speedups while maintaining modest memory usage and strong parallel scalability. As genome annotations continue to grow in complexity and size, *GFFx* offers a practical and extensible foundation for future bioinformatics workflows. While robust for standard GFF3 files, the current implementation assumes well-formed input and does not yet support GTF or legacy GFF2 formats. Enhancing compatibility and fault tolerance— particularly for nonstandard annotations—remains an important area for development. Planned extensions include support for additional formats, distributed computing integration, and interactive search for large-scale databases. *GFFx* is distributed as a statically compiled binary for Linux, macOS, and Windows. It can also be used as a Rust library, allowing integration into custom pipelines and tools. Its modular architecture and clean API offer fine-grained access to core functions, making *GFFx* both performant and programmable. Full documentation is available at <u>docs.rs/GFFx</u>, and the GitHub repository includes user manuals, benchmarks, input data, and source code for complete reproducibility (https://github.com/Baohua-Chen/GFFx).

## Methods

### Architectural design of indexing system underpins GFFx performance

*GFFx* was developed as a modular and high-performance command-line toolkit for processing large GFF files. Its efficiency is supported by a carefully engineered indexing system (**Fig. 1**). At the core of *GFFx* are two index files shared across all subcommands: .*prt* and .*gof*. The .*prt* index encodes the hierarchical relationships among annotated features and delineates annotation blocks as minimal, biologically coherent units, such as complete gene models or transcript structures. The .*gof* index maps each annotation block to its corresponding byte-offset range in the original GFF file, enabling direct memory-mapped access to specific regions without requiring full-file scanning or decompression. Together, these two indices provide the structural and positional backbone of *GFFx*, allowing fast and model-aware access to genome annotations with minimal input/output overhead. To minimize redundancy and reduce index file size, both .*prt* and .*gof* use numeric feature identifiers assigned in order of appearance. The original string-form feature IDs are stored separately in the .*fts* file.

In addition to the core indices, *GFFx* generates several auxiliary index files that support specific subcommands. The extract subcommand retrieves the full annotation block associated with a given feature and requires only the .*fts* index, which records all feature identifiers in order, together with the .*prt* and .*gof* files. For attribute-based queries, the .*atn* file stores all user-specified string-form identifiers found in the attribute field of the GFF file (such as “gene”, “Name”, or “symbol”), while the .*a2f* file maps each attribute value to its corresponding numeric feature ID. These two files are used by the search subcommand, which enables both exact and fuzzy attribute queries. The intersect subcommand uses an interval tree scheme. *GFFx* builds a .*rit* file containing all interval-tree nodes laid out sequentially and a companion .*rix* file that records offsets in .*rit* for each chromosome or scaffold, so that only the relevant subtree is loaded on demand. This reduces region-query time complexity from O(N) to O(log N), greatly speeding up lookups in large genomes. All indices are written in compact binary format and accessed on demand by each subcommand to minimize storage footprint and loading time.

### Efficient Runtime Strategies for Feature and Region-Based Extraction

To achieve high-throughput querying from ultra-large GFF3 files, *GFFx* incorporates several performance-oriented design strategies beyond its indexing system. All subcommands operate directly on memory-mapped representations of the original GFF file using the memmap2 library. This eliminates the need for repeated I/O or line-by-line parsing by allowing byte-range access to annotation blocks through read-only mappings. Extracted regions or feature models are located via index lookups and retrieved efficiently by copying their byte slices directly from the memory-mapped buffer. To minimize redundant computation, *GFFx* leverages reference-counted shared memory to ensure that index structures such as .*gof* and .*rit* are loaded only once and reused across all operations. Output blocks are streamed directly to disk, avoiding large memory buffers, and the software assumes well-formed GFF3 input to reduce validation overhead.

To ensure high-performance region-based feature extraction, *GFFx* leverages several optimizations provided by the Rust ecosystem, such as the use of “FxHashMap” for low-overhead hash-based mappings and “lexical_core” for converting ASCII byte sequences into integer coordinates with minimal latency. Additionally, input regions are pre-bucketed by chromosome and sorted by the start coordinates, ensuring each interval tree to be queried only with relevant regions, thereby reducing unnecessary computation and improving cache locality.

## Supplementary data

Supplementary data are available online.

## Data availability

The source code and a comprehensive user manual are available on GitHub: https://github.com/Baohua-Chen/GFFx. Benchmarking scripts and original results are also publicly available at: https://github.com/Baohua-Chen/GFFx_benchmarks.

## References

1. Moore JE, Purcaro MJ, Pratt HE, Epstein CB, Shoresh N, Adrian J, et al. Expanded encyclopaedias of DNA elements in the human and mouse genomes. Nature. Nature Publishing Group; 2020; doi: 10.1038/s41586-020-2493-4.

2. Forrest ARR, Kawaji H, Rehli M, Kenneth Baillie J, de Hoon MJL, Haberle V, et al. A promoter-level mammalian expression atlas. Nature. Nature Publishing Group; 2014; doi: 10.1038/nature13182.

3. Satterlee JS, Chadwick LH, Tyson FL, McAllister K, Beaver J, Birnbaum L, et al. The NIH Common Fund/Roadmap Epigenomics Program: Successes of a comprehensive consortium. Science Advances. American Association for the Advancement of Science; 2019; doi: 10.1126/sciadv.aaw6507.

4. Harrison PW, Amode MR, Austine-Orimoloye O, Azov AG, Barba M, Barnes I, et al. Ensembl 2024. Nucleic Acids Research. 2024; doi: 10.1093/nar/gkad1049.

5. Jeffrey M. Perkel. Why scientists are turning to Rust. Nature. 588:1852020;

6. Köster J. Rust-Bio: a fast and safe bioinformatics library. Bioinformatics. 2016; doi: 10.1093/bioinformatics/btv573.

7. Huey JD, Abdennur N. Bigtools: a high-performance BigWig and BigBed library in Rust. Bioinformatics. 2024; doi: 10.1093/bioinformatics/btae350.

8. Vijendran S, Anderson T, Markin A, Eulenstein O. Phylo-rs: an extensible phylogenetic analysis library in rust. BMC Bioinformatics. 2025; doi: 10.1186/s12859-025-06234-w.

9. Jha A, Bohaczuk SC, Mao Y, Ranchalis J, Mallory BJ, Min AT, et al. DNA-m6A calling and integrated long-read epigenetic and genetic analysis with fibertools. Genome Res. 2024; doi: 10.1101/gr.279095.124.

10. Dale R. Gffutils: GFF and GTF file manipulation and intercon-version. Github;

11. Pertea G, Pertea M. GFF Utilities: GffRead and GffCompare. F1000Res. 2020; doi: 10.12688/f1000research.23297.2.

12. Chapman B. bcbio-gff. GitHub. v0. 6.4.

13. Dainat J. Another Gtf/Gff Analysis Toolkit (AGAT): Resolve Interoperability Issues and Accomplish More with Your Annotations. Plant and Animal Genome XXIX Conference (January 8-12, 2022). PAG;

14. Quinlan AR, Hall IM. BEDTools: a flexible suite of utilities for comparing genomic features. Bioinformatics. 2010; doi: 10.1093/bioinformatics/btq033.

15. Lawrence M, Huber W, Pagès H, Aboyoun P, Carlson M, Gentleman R, et al. Software for Computing and Annotating Genomic Ranges. PLOS Computational Biology. Public Library of Science; 2013; doi: 10.1371/journal.pcbi.1003118.

