## Supplementary Fig. for "GFFx: A Rust-based suite of utilities for ultra-fast genomic feature extraction"

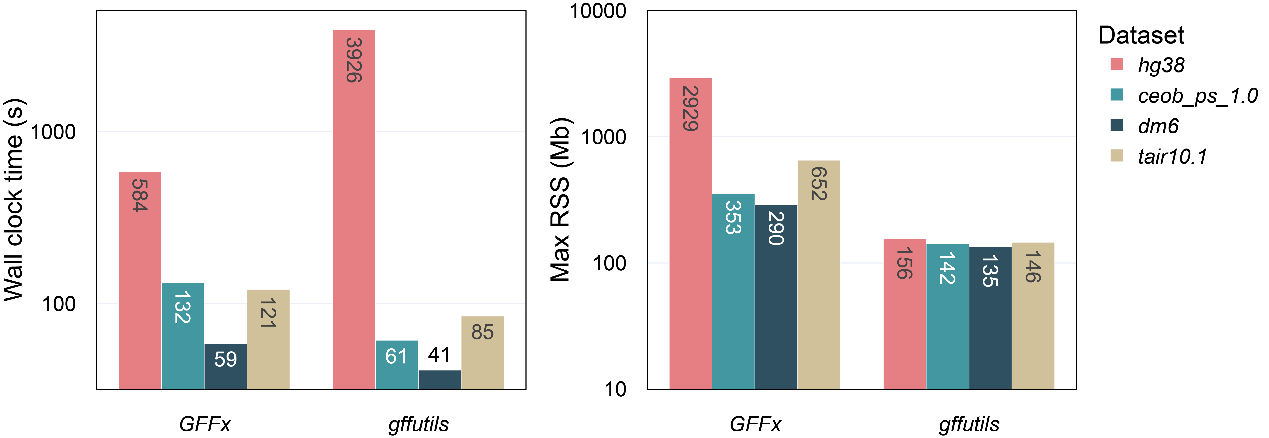

**Figure S1. Comparison of preprocessing performance between *GFFx* and *gfftuils*.** (a) Median wall-clock time (log scale) on *hg38* (pink), *ceob_ps_1.0* (cyan), *dm6* (dark teal), and *tair10.1* (tan) using *GFFx* and *gffutils*. (b) Maximum resident set size (RSS, log scale), a measure of peak memory consumption, for each tool and dataset.
